# Single-vector AAV delivery of En-ISPmu1 and donor DNA for *in vivo* targeted genome integration

**DOI:** 10.64898/2026.10.26.761871

**Authors:** Alessio Conci, Ilaria Bonuzzi, Matteo Ciciani, Federica Esposito, Pasquale Piccolo, Fanny Collaud, Giuseppe Ronzitti, Alberto Auricchio, Nicola Segata, Anna Cereseto

## Abstract

TnpBs are compact RNA-guided nucleases ideally suited for *in vivo* delivery. By screening novel TnpBs identified from large metagenomic datasets, ISPmu1 was discovered and its protein and ωRNA were engineered to generate En-ISPmu1, which achieved editing efficiencies exceeding 65% across 12 human loci. Owing to its compact size, En-ISPmu1 was delivered together with a donor DNA in a single AAV vector, yielding up to 12.7% targeted integration in mouse liver. These findings establish En-ISPmu1 as a promising versatile platform for *in vivo* therapeutic genome editing applications.

## Main text

Inherited disorders often arise from a broad spectrum of pathogenic variants. Although the precision and versatility of modern genome editing technologies have created new opportunities for personalized medicine, allelic heterogeneity remains a major barrier to therapeutic development. Designing, validating, and manufacturing a bespoke treatment for every disease-causing variant limit the long-term clinical scalability. Targeted insertion of a functional cDNA into a safe-harbor site^1^ or the endogenous disease-associated locus^2^ could provide a universal gene therapy strategy, restoring gene function across diverse pathogenic variants in patients with the same genetic disorder. Adeno-associated viral (AAV) vectors are currently among the most common and efficient delivery platforms for *in vivo* gene therapy^3^; however, their application is constrained by their limited cargo capacity. To establish a platform for site-specific integration, we searched for miniature RNA-guided nucleases suitable to be packaged together with a donor DNA in an all-in-one AAV vector was performed. We focused on TnpB nucleases, encoded by IS200/605 insertion sequences (IS)^4^, due to their compact size (around 400 aa)^5,6^.

To explore TnpB diversity and identify novel active variants, we screened a large metagenomic database comprising 156,734 isolate genomes and 864,853 metagenome-assembled genomes (MAGs)^7,8^. We identified 337,477 putative TnpB orthologs, revealing extensive diversity across Bacteria and Archaea (**Extended Data Fig. 1**). To select TnpB variants for genome editing in human cells, we first analyzed known variants active in *E. coli*^9^ and identified 20 highly conserved amino acid residues (**Supplementary Fig. 1**), which were subsequently used to filter potential candidates. Next, by multiple sequence alignment we identified the boundaries of the newly identified TnpB-encoding IS elements, generating *in silico* predictions of the cognate ⍰RNA and transposon-adjacent motif (TAM) sequences (**Fig. 1a**). TAMs were identified from the 4-5 most conserved nucleotides at the IS 5’-end^5,9,10^ and candidate ⍰RNA sequences starting 220 bp upstream of the transposon-encoded motif (TEM) at the IS 3’-end. For each TnpB-⍰RNA pair, we predicted the ⍰RNA secondary and ternary complex structures, and candidates with poor predicted folding were removed. In total, we selected 25 phylogenetically diverse TnpB orthologs for experimental characterization (**Fig. 1b** and **Supplementary Tab. 1**).

**Figure 1.**
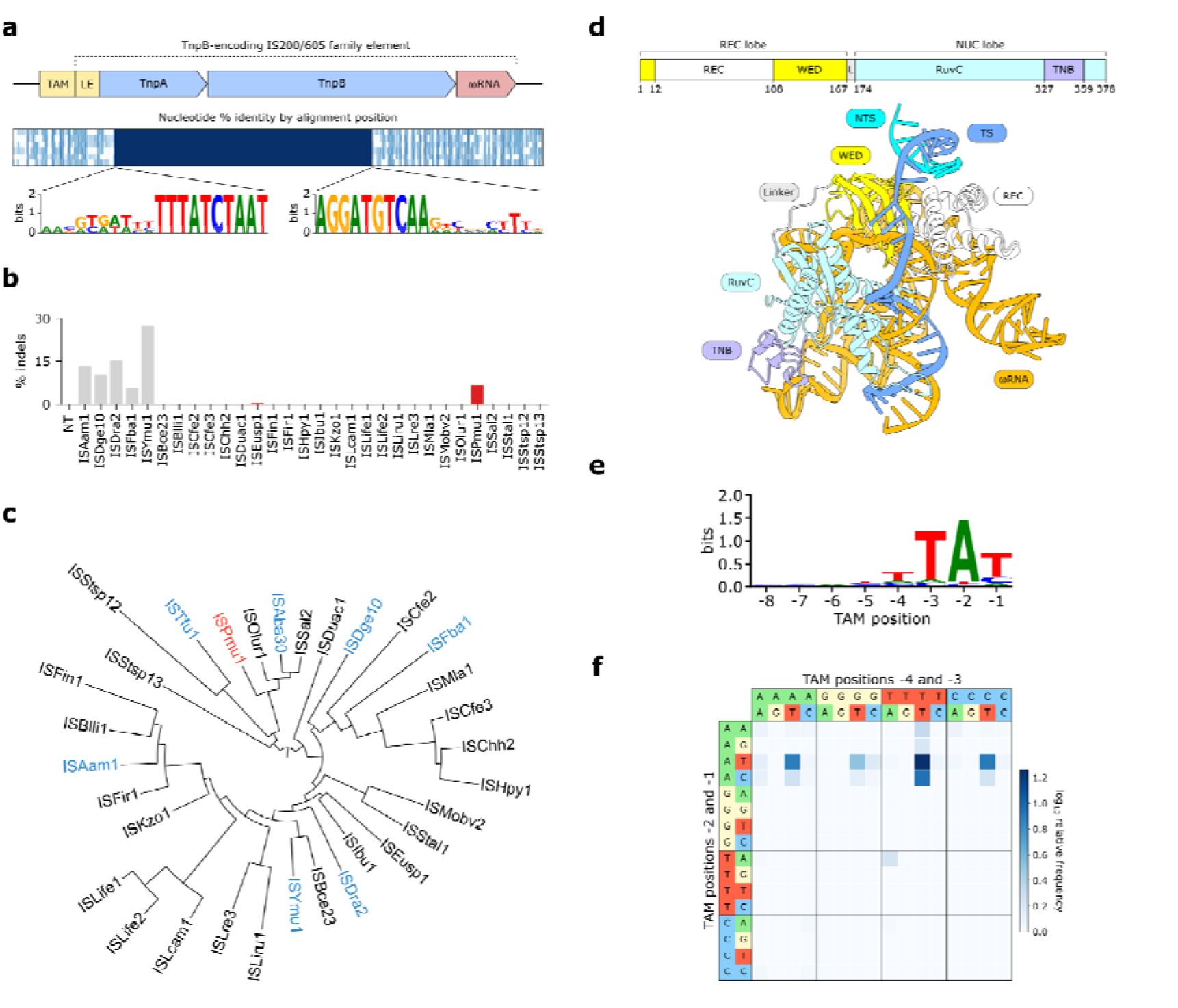
Identification of ISPmu1 and its TAM. (**a**) TAM and ⍰RNA sequence prediction from the 5’ and 3’ ends of IS200/605 elements encoding TnpB candidates. (**b**) Maximum likelihood phylogenetic tree of 25 TnpBs selected for experimental characterization (black) and 7 known TnpBs active in human cells (blue). ISPmu1 is indicated in red. (**c**) Indels generated at the *EMX1* locus by TnpB candidates (red) and positive controls (gray). The best-performing guide for each ortholog is shown. (**d**) ISPmu1 predicted domain architecture and ternary complex structure. (**e**) Sequence logo of ISPmu1 TAM. (**f**) TAM heatmap quantifying the relative enrichment of each cleaved nucleotide combination.

To directly test candidate variants for activity in human cells, we co-transfected plasmids expressing human codon-optimized TnpB candidates with cognate ⍰RNAs in HEK293T cells and indel formation was assessed by deep sequencing at a benchmark locus, *EMX1*^5,9^. One active variant was identified, namely ISPmu1 (378 aa) from *Pasteurella multocida* (**Fig. 1c**), with an editing efficiency of 6.5% (**Fig. 1d** and **Supplementary Fig. 2**). We experimentally validated the *in silico* predicted TAM of ISPmu1, revealing preferential cleavage of target sequences flanked by a 5’-TTAT-3’ motif (**Fig. 1e-f** and **Supplementary Fig. 3**).

To improve ISPmu1 editing efficiency, we optimized both the ωRNA scaffold and the TnpB protein. Since ⍰RNA shortening has been reported to enhance TnpB activity^11–13^, we tested several 5’-truncated ⍰RNAs, including one rationally designed (⍰RNA-136 nt) based on secondary structure prediction (**Supplementary Tab. 2**). Notably, this variant showed significantly higher activity than the original 220 nt ⍰RNA (**Fig. 2a**). The trimmed ⍰RNA scaffold was further engineered following secondary structure and ternary complex predictions, thereby identifying modifiable stem-loops (**Supplementary Tab. 3**), resulting in ⍰RNA-v12 and ⍰RNA-v14 with increased activity by 1.3- and 1.5-fold, respectively (**Fig. 2b** and **Extended Data Fig. 2**). By combining modifications from ⍰RNA-v12 and ⍰RNA-v14 we generated a shorter ⍰RNA (117+G nt), which maintained a robust editing activity while providing a further compact configuration (**Fig. 2c** and **Extended Data Fig. 3**).

**Figure 2.**
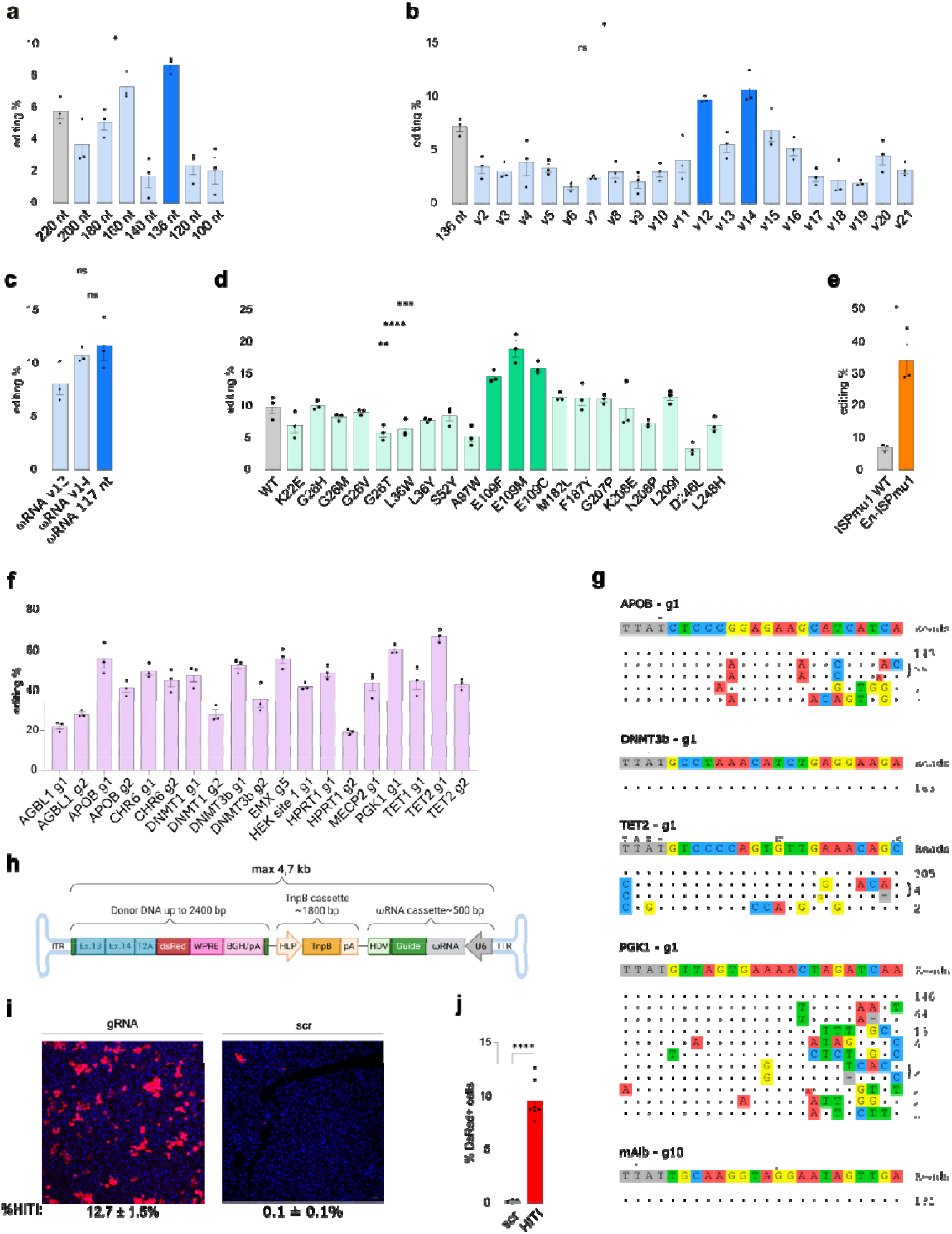
ISPmu1 optimization, characterization, and *in vivo* application. (**a**) ISPmu1 editing efficiencies using ωRNAs of various lengths (nt). (**b**) Editing efficiency of rationally designed ωRNAs (**Extended Data Fig. 2**). (**c**) Editing efficiency of best-performing rationally designed ωRNA variants (v12 and v14) and their combination into ωRNA 117 nt. (**d**) Editing activity of ISPmu1 single amino acid mutants at the *EMX1* genomic locus. (**e**) Editing efficiencies of ISPmu1 and En-ISPmu1 at the *EMX1* locus. (**f**) Deep sequencing analysis of En-ISPmu1 indels at 12 human endogenous loci. (**g**) GUIDE-seq analysis at the indicated loci. (**h**) Schematic representation of the AAV8 vector used for HITI *in vivo* experiments. (**i**) Representative fluorescence microscopy images of liver cryo-sections from wild-type mice injected with AAV-HITI and relative percentage of DsRed+ cells. (**j**) Quantification of integrated DsRed from fluorescence microscopy images. Data are represented as mean ± SEM of n=3 independent replicates (**a**, **b**, **c**, **d**, **e**, **f**) and n=7 independent mouse liver samples (**j**). Comparisons for panels **a**, **b**, **c,** and **d** were performed using a one-way ANOVA followed by Dunnett’s multiple comparisons test against the references; comparisons for panels **e** and **j** were performed using an unpaired t-test with Welch’s correction. Statistical significance for adjusted p-values was defined as: ns p ≥ 0.05, * p < 0.05, ** p < 0.01, *** p < 0.001, and **** p < 0.0001.

Screening for guide length revealed 20 nt as the optimal size (**Supplementary Tab. 4** and **Supplementary Fig. 4**). To optimize ISPmu1, we tested various nuclear localization signals (NLS) and tags combinations (**Supplementary Fig. 5**)^12^, leading to the identification of architecture 2 (ARC-2) as the optimal configuration for its compact size (438 aa) and editing efficiency (**Supplementary Fig. 6**). To further enhance ISPmu1 activity, we tested 18 substitutions orthologous to mutations that improved ISDra2 TnpB performance^13^ and subsequently identified three E109 substitutions (C, F and M) that significantly increased editing activity (**Fig. 2d**).

The combination of ISPmu1-E109M as ARC2 together with the 117nt ⍰RNA, hereafter designated En-ISPmu1, achieved a 5-fold increase in activity over the original system (**Fig. 2e**) and was then evaluated across a panel of 12 endogenous human loci via deep sequencing. Editing activity was detected at all tested loci with efficiencies ranging from 19.2% to 67.3% and averaging over 40%, demonstrating that En-ISPmu1 has a broad targeting range and robust activity (**Fig. 2f** and **Supplementary Fig. 7**). We tested genome-wide specificity through GUIDE-seq^14^ across the top 4 most efficient guides from **Figure 2f**^14^. While *DNMT3b* and *TET2* loci showed no or low off-targets, *APOB1* and *PGK1* editing was associated with higher non-specific cleavages, indicating variable specificity throughout the genome (**Fig. 2g**).

Beyond nuclease activity, En-ISPmu1 can be used as a base editor by eliminating its nuclease activity (D181A RuvC mutation) and fusing it with the adenine deaminase Tad8e at the N-terminus. dEn-ISPmu1-Tad8e displayed an editing window ranging from position 4 to 12 of the target sites, with an A-T to G-C conversion of up to 30% (**Extended Data Fig. 4**).

Next, we leveraged the compact size of En-ISPmu1 to evaluate *in vivo* homology-independent targeted integration (HITI) using an all-in-one AAV vector that simultaneously delivered the RNA-guided nuclease and donor DNA (**Fig. 2h**)^15^. We selected the highly expressed hepatic albumin locus as a well-established site for therapeutic gene insertion in the treatment of metabolic diseases^2,16,17^. A construct designed to include the DsRed expression marker alongside the TnpB and guide RNA was packaged in a single AAV8 vector; the targeting guide was selected among those with higher efficiency and specificity (**Fig. 2h** and **Extended Data Fig. 5**). The HITI donor construct contained the last mouse albumin exons (13 and 14) in frame with the promoter-less coding sequence of the DsRed fluorescent reporter transgene, enabling direct visualization of edited hepatocytes. Neonatal wild-type mice were injected via the temporal vein, and liver tissues were analyzed one month later via confocal microscopy to quantify DsRed-positive (DsRed+) hepatocytes. Quantification showed a mean of 9.5% DsRed+ cells across all mice, with robust and uniform expression throughout the liver (**Fig. 2i-j** and **Extended Data Fig. 6**). We confirmed the correct integration of the target sequence by PCR of genome-donor DNA junctions (**Supplementary Fig. 8**).

In summary, we developed a computational discovery pipeline enabling the identification of ISPmu1 from an extensive metagenomic dataset. Engineering of its protein and ωRNA increased the activity by 5-fold, yielding up to 67% editing at endogenous human loci. The compact size and robust editing efficacy of the resulting En-ISPmu1 permitted *in vivo* targeted integration through a single AAV vector. Together, these findings establish a versatile genome editing platform and provide a strategy for overcoming cargo limitations of AAV-based gene therapy.

## Data availability

Sequencing data for TAM determination assays, GUIDE-seq, and deep sequencing experiments are publicly available at NCBI Sequence Read Archive (PRJNA1533856).

## Code availability

Custom code used in this study is publicly available at https://github.com/Matteo-Ciciani/TnpB-data-analysis.

## Acknowledgments

We are grateful to Cereseto’s lab members for helpful discussions throughout the project. We thank the Next Generation Sequencing facility at the University of Trento for technical support. This work was supported by the Horizon Europe EIC Pathfinder programme AAVolution (grant agreement 101071041) to A.Ce. and A.A., by Fondo Italiano per la Scienza (FIS) from the Italian Ministry for Research (MyBET grant agreement FIS00002542) to A.Ce., by the European Research Council (ERC-CoG microTOUCH-101045015) to N.S, and the ERC Advanced EXPEDITE grant (grant number 101097155).

## Author information

A.Co, I.B. and F.E. designed and performed the experiments; M.C. developed the computational pipeline for TnpB identification; A.Co, I.B, M.C., F.C. and F.E. collected and analyzed the data; A.Ce., N.S., A.A., P.P., G.R., A.Co, I.B, M.C., F.C. and F.E. conceived and designed the study. A.Co, I.B, M.C., and A.Ce wrote and edited the paper; A.Ce coordinated the study. All authors read, corrected, and approved the final manuscript.

## Ethics declaration

All animal procedures were performed in accordance with the Italian Ministry of Health regulations governing the use of animals for scientific purposes and under the approved experimental protocol (Ministry of Health authorization no. 625/2023-PR).

## Methods

### Identification of TnpB proteins

A catalog of bacterial and archaeal genomic sequences was retrieved from 156,734 publicly available isolate sequences from the NCBI database^1^ (available as of January 2023) and 864,853 metagenome-assembled genomes (MAGs) from Blanco-Míguez, A. *et al.* 2023^2^. Proteins were predicted for all genomes using Prokka (v1.14)^3^. TnpB profile HMMs were retrieved from the NCBI CDD database (accession numbers NF038280, NF038281, NF040570) and searched against all proteins using hmmsearch v3.3.2 with parameters -E1^4^. A total of 402,420 matching proteins between 300 and 600 aa were retrieved. To remove truncated sequences, we clustered proteins at 70% identity and 95% coverage with mmseqs cluster v2.13 with parameters -c 0.95 --cov-mode 0 --min-seq-id 0.7 --cluster-reassign^5^. For each cluster, we built a multiple sequence alignment using MAFFT v7.490 --maxiterate 50 -- op 5.5 --ep 1.5^6^, which was used to generate an HMM profile that was aligned with the TnpB models used above with HHAlign v3.3.0^7^. Matches scoring below 50 were discarded, and query and template alignment coverage was computed. Matches with both alignment coverages below 0.3 or with query coverage > 0.9 and template coverage < 0.75 (i.e., truncated alignments) were discarded. The remaining 346,041 sequences were reclustered at 95% identity and 60% coverage with mmseqs cluster v2.13 with parameters -c 0.6 --cov-mode 5 --min-seq-id 0.95 --cluster-reassign to further remove truncated proteins. For each cluster, sequences with length below 90% of the cluster median length were discarded, retaining a total of 337,477 putative TnpBs.

### Identification of Cas12 proteins

CRISPRCasTyper v1.8.0 was run on all genomes to identify CRISPR arrays and Cas proteins^8^. To distinguish between TnpB and small Cas12 variants, for each putative TnpB locus, we annotated the presence of a Cas1 or Cas2 protein or a CRISPR array within a 3,000 bp flanking region. In total, 6,582 proteins were reclassified as putative Cas12 and were not considered for candidate TnpB selection.

### TnpB phylogenetic tree

The 337,477 sequences previously identified, together with 21,878 Cas12 proteins identified with CRISPRCasTyper, were dereplicated at 98% identity and coverage using mmseqs cluster v2.16 with parameters -c 0.98 --cov-mode 0 --min-seq-id 0.98 --cluster-reassign. To further remove truncated Cas12 sequences, all dereplicated sequences were clustered at 70% identity and coverage using mmseqs cluster v2.16 with parameters -s 7 -c 0.7 --cov-mode 1 --cluster-mode 2 --min-seq-id 0.7. For each cluster, sequences below 80% of max cluster sequence length were discarded. The remaining 64,316 dereplicated sequences were clustered at 50% identity and 70% coverage using mmseqs cluster v2.16 with parameters -s 7 -c 0.7 --cov-mode 0 --min-seq-id 0.5 --cluster-reassign, obtaining 4,018 clusters. For each cluster, the longest sequence was picked as representative. Sequences were aligned using MAFFT 7.525 --maxiterate 100. The alignment was trimmed using trimal^9^ v1.4 -gt 0.1, and 64 sequences composed of more than 50% gaps were removed. An initial tree was built using IQ-TREE version 2.0.3 with parameters -B 1000 and automatic model selection^10^. The alignment was sorted using the maximum likelihood tree and visually inspected, revealing several fragmented sequences that escaped previous filtering steps. To improve the accuracy of the alignment, key regions corresponding to the N-terminal region, the ZF domain, and the RuvC-I, -II, and -III domains were extracted from the alignment. For each sequence, gap fraction was computed for all key regions, and sequences with high gap fraction within that region were discarded unless they belonged to a clade of 5 or more sequences with conserved region deletion. A new tree was built with the 3,330 remaining sequences using FastTree v2.1.11 -spr 4 -slownni -mlacc 2. To mitigate long branch attraction, leaf branches longer than 1.5 and internal branches longer than 1 were identified and pruned if they contained fewer than 5 sequences, to avoid eliminating highly divergent clades. This process was iterated three times, removing a total of 35 sequences. The final tree was built using IQ-TREE version 2.0.3 with parameters -B 1000 --ntop 100 --nbest 20 and automatic model selection. The tree was visualized using GraPhlAn version 1.1.3^11^.

### Selection of TnpB variants

To select candidate TnpB variants, the number of copies per genome was computed by clustering them at 99% identity and coverage using mmseqs cluster v2.16 with parameters - c 0.99 --cov-mode 0 --min-seq-id 0.99 --cluster-reassign. 398 TnpBs with 5 or more copies per genome were then compared with variants already characterized in previous studies^12,13^ by clustering them at 90% identity and 80% coverage using mmseqs cluster v2.16 with parameters -c 0.8 --cov-mode 0 --min-seq-id 0.9 --cluster-reassign. TnpB candidates that co-clustered with previously characterized variants were removed. To identify residues conserved in active variants, proteins that showed activity in *E. coli*^12^ were aligned with MUSCLE v5.1^14^. The alignment was trimmed using trimal v1.4 -gappyout and used to build an HMM profile with hmmbuild v3.3.2, which was visualized using hmmlogo v3.3.2. A set of 20 residues conserved in >90% of active variants was identified, of which 10 were already reported by Xiang, G. *et al.* (2023)^12^. TnpB candidates were aligned to active variants in Xiang, G. *et al.* (2023) ^12^ using MAFFT v7.525 --maxiterate 100, and variants lacking more than 2 conserved residues were discarded. The remaining variants were further analyzed to predict TAM and RNA sequences. For each variant, 5000 bp sequences flanking every TnpB copy were extracted from genomes and aligned with MAFFT v7.525 --maxiterate 10. The alignment was analyzed to identify conserved transposable element ends: TAM predictions were generated by identifying the first 4 or 5 conserved nucleotides at the 5’ end, while RNAs were predicted as sequences starting 220 bp upstream from the 3’ end. The RNA secondary structure was inspected using mfold^15^, while the TnpB structure was inspected by generating ternary complex structure predictions using AlphaFold3^16^. In total, 25 TnpB variants with clearly predicted TAM and RNA sequences were selected for experimental characterization.

### *In vitro* TAM assay

The *in vitro* TAM determination assay was performed by adapting the protocol from Karvelis et al. 17. The ISPmu1 human codon-optimized DNA sequence was cloned into the expression vector pT7-N-His-GST (Thermo Fisher Scientific) for in vitro transcription and translation (IVT) as described above. The IVT reaction was performed using the 1-Step Human High-Yield Mini IVT Kit (Thermo Fisher Scientific) according to the manufacturer’s protocol. A DNA fragment containing the T7 promoter and ISPmu1 ⍰RNA sequence was synthesized as complementary oligonucleotides, annealed, and then in vitro transcribed using the HighYield T7 RNA Synthesis Kit (Jena Bioscience), following the manufacturer’s guidelines. Purification of the in vitro transcription reaction was done by overnight ammonium acetate precipitation. The ISPmu1-⍰RNA ribonucleoprotein (RNP) complex was assembled by combining 20 μL of the supernatant containing soluble ISPmu1 protein with 1 μL of RiboLock RNase Inhibitor (Thermo Fisher Scientific) and 2 μg of ⍰RNA. The RNP was incubated at room temperature for 30 min. The RNP was subsequently incubated with 1 μg of 7N randomized TAM plasmid DNA library 18 and cleavage buffer (10 mM Tris-HCl (pH 7.5 at 37 °C), 100 mM NaCl, 1 mM DTT, and 10 mM MgCl2)19 at 37 °C for 3 h. The ends of the cleaved TAM library were repaired with 1 μL T4 DNA polymerase and dNTPs; 3’-dA overhangs were added with 1 μL DreamTaq, and RNA was removed with RNase A/T1. The final product was purified using a GeneJet PCR Purification Kit (Thermo Fisher Scientific). A double-stranded DNA adapter was ligated to the TAM library DNA ends, and the final ligation product was purified with AMPure XP beads (Beckman Coulter) at a 1:0.8 ratio. A first PCR (Phusion HF DNA polymerase, Thermo Fisher Scientific) was performed to enrich the cleaved sequences using a forward primer annealing to the adapter and a reverse primer designed on the plasmid backbone downstream of the TAM 18. A second round of PCR was performed to attach Illumina indexes and adapters. PCR products, in both steps, were purified using AMPure XP beads (Beckman Coulter) (ratio 1:1). The library was analyzed by 150 bp single-end reads sequencing on an Illumina MiSeq sequencer. TAM sequences were extracted from Illumina MiSeq reads and used to generate TAM sequence logos, using Logomaker version 0.8 20. TAM heatmaps 21 were used to display TAM enrichment, computed as the frequency of TAM sequences derived from target cleavage events divided by the background frequency of uncleaved sequences.

### Mammalian expression plasmids and AAV vector constructs

A pcDNA3-derived plasmid was used to express the selected TnpBs in mammalian cells, while the guides with their ωRNAs were expressed using pUC-derived plasmids. Briefly, pcDNA3 was modified by substituting the SpCas9 and its sgRNA scaffold with the human codon-optimized sequence of the TnpB orthologs. The TnpB coding sequences and the ωRNA scaffolds were obtained as synthetic fragments from either Genscript or Twist Bioscience; all TnpB candidates were fused with a V5 tag and an SV40 NLS at the N-terminus and a nucleoplasmin nuclear localization signal followed by a 3xHA-tag at the C-terminus. Optimization steps involved changing the NLS architecture by testing the following configurations^22^: ARC-1 (SRAD+SV40 at C-terminus) and ARC-2 (SRAD+SV40+GS-linker+BPNLS). Spacer sequences were cloned into the pUC plasmids as annealed DNA oligonucleotides containing a variable 10-20-nt spacer sequence using double BbsI sites present in the plasmid. The list of spacer sequences used is reported in **Supplementary Table 5**. The plasmid used for AAV vector production was kindly gifted by Alberto Auricchio at TIGEM (Naples). The plasmid contained the inverted terminal repeats of AAV serotype 2, and the scrambled guide was designed so as not to align with any sequences in the mouse genome.

### Cell lines and culture conditions

HEK293T cells (obtained from ATCC, #CRL-11268) and HEPA cells (kindly gifted by Pasquale Piccolo at TIGEM, Naples) were cultured in DMEM (Life Technologies) supplemented with 10% FBS (Life Technologies), 2 mM GlutaMax (Life Technologies), and penicillin/streptomycin (Life Technologies). All cells were incubated at 37 °C and 5% CO_2_ in a humidified atmosphere. All cells tested mycoplasma negative (PlasmoTest, Invivogen).

### Cell line transfections

For editing analyses of genomic loci, 150,000 HEK293T cells were seeded in a 24-well plate 24 h before transfection. Cells were then transfected with 600 ng of pcDNA3-TnpB plasmid and 400 ng of a pUC-ωRNA plasmid, using the TransIT-X2 reagent (Mirus Bio) according to the manufacturer’s protocol. Cell pellets were collected 3 days post-transfection for indel evaluations.

### Evaluation of editing efficiency

DNA was extracted using the DNeasy Blood & Tissue kit or QuickExtract DNA Extraction Solution (Lucigen) for cell culture 3 days after transfection. Editing efficiency of TnpB candidates and of ISPmu1 on human and mouse endogenous loci was analyzed by deep sequencing, amplifying genomic sites of interest with primers containing Illumina forward and reverse adaptor sequences and using Phusion HF DNA polymerase (Thermo Fisher Scientific), using as template 400 ng of DNA (from DNeasy Blood & Tissue kit) or 2 μl from the QuickExtract DNA Extraction Solution. The oligonucleotides used are listed in **Supplementary Table 6**. The amplified products were purified (Agencourt AMPure beads), quantified with the Qubit, and used for a second round of PCR (Phusion HF DNA polymerase - Thermo Fisher Scientific) to attach the Illumina indexes. After final purification and quantification, a pool of each sample at the same nM was analyzed with 250 bp paired-end sequencing, using a v2 flow cell on an Illumina MiSeq sequencer. Editing levels were assessed using CRISPResso2^23^ v2.3.1 with parameters --default_min_aln_score 10 -- min_bp_quality_or_N 20 --ignore_substitutions, with a 40 bp quantification window centered on the guide 3’ end, running in mixed pooled mode and aligning reads to the GRCh38.p14 human genome assembly. To amplify the target locus EMX1 for testing the ωRNA modifications, ISPmu1 mutations, and guide length experiments, PCR reactions were performed using the HOT FIREPol DNA polymerase (Solis BioDyne) and the oligonucleotides listed in **Supplementary Table 5**. The amplified products were purified, Sanger sequenced (Microsynth) and analyzed with the TIDE web tool version 3.30 (http://shinyapps.datacurators.nl/tide/).

### Guide-seq

GUIDE-seq experiments were performed as described in previous studies^24^. Briefly, HEK293T and HEPA cells were transfected using Lipofectamine 3000 transfection reagent (Invitrogen) with 1 μg (600 ng TnpB plasmid + 400 ng ωRNA plasmid) and 2 μg (1200 ng TnpB plasmid + 800 ng ωRNA plasmid) of plasmid, respectively, and 10 pmol of dsODNs. Three days after transfection, cells were collected, and genomic DNA was extracted using NucleoSpin Tissue Mini kit (Macherey-Nagel) following the manufacturer’s instructions. Genomic DNA was sheared to an average length of 500 bp using a focused ultrasonicator (Covaris). An end-repair reaction was performed using NEBNext Ultra End Repair/dA Tailing Module and adaptor ligation using NEBNext Ultra Ligation Module, as described by Nobles *et al.*^25^. Amplification steps were then performed following the GUIDE-seq protocol from Tsai *et al.*^24^. Following quantification of the libraries by Qubit dsDNA High Sensitivity Assay kit (Invitrogen), MiSeq sequencing system (Illumina) was used with an Illumina MiSeq Reagent kit V2-150PE. Raw sequencing data (FASTQ files) were analyzed using the GUIDE-seq computational pipeline available at https://github.com/tsailabSJ/guideseq.

### AAV vector production and characterization

AAV8 vectors were selected for *in vivo* hepatic delivery based on their high liver tropism and efficient transduction of hepatocytes^26^. Vectors were produced by InnovaVector S.r.l. through triple transfection of HEK293 cells, followed by two rounds of CsCl₂-based purification, according to previously described procedures^27^. For each vector preparation, the viral titer, expressed as genome copies per milliliter (gc/mL), was determined by averaging the values obtained from three independent rounds of TaqMan quantitative PCR.

### Systemic vector administration

Neonatal C57BL/6J mice received 1.2 × 10^14 genome copies (GC)/kg by temporal vein injection on postnatal days (P) 1–2, as previously described by Gombash Lampe *et al.*^28^.

### Liver fluorescence imaging

To evaluate HITI-mediated integration in the liver, mice were euthanized at postnatal day 30 (P30), and livers were collected as previously reported^26^. Briefly, small fragments from each liver lobe were fixed overnight in 4% PFA, cryoprotected in 15% sucrose for 24 h followed by 30% sucrose overnight, embedded in O.C.T. compound (Cat# 0782, Kaltek), and cryosectioned at 5 μm. Sections were mounted with Vectashield containing DAPI (Cat# H-1200-10, Vector Laboratories). Fluorescence imaging was performed using the Nikon Ti2 Eclipse confocal microscope system (Nikon Corporation, Tokyo, Japan) with appropriate excitation and emission settings for DsRed and DAPI. Three to four representative fields were acquired from each liver at 20× magnification and analyzed using ImageJ/Fiji software (https://imagej.net/). DsRed-positive hepatocytes, indicative of HITI-mediated integration, were identified based on their characteristic morphology and quantified relative to the total number of DAPI-positive nuclei. HITI efficiency was calculated as the percentage of DsRed-positive hepatocytes over the total DAPI-positive cells.

## EXTENDED DATA

**Extended Data Fig. 1.**
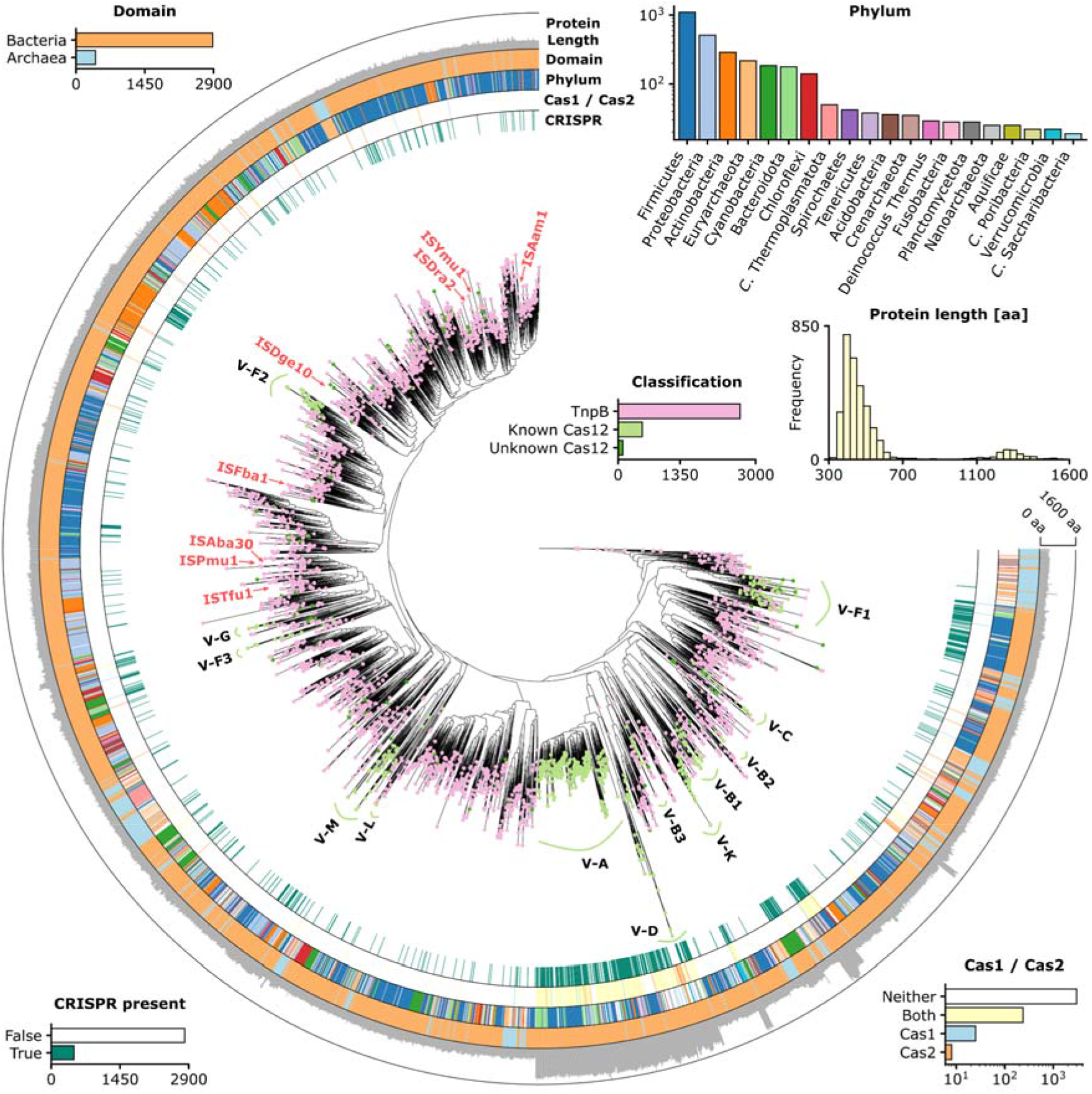
TnpB phylogenetic tree. Maximum-likelihood phylogenetic tree of 3,295 TnpB and Cas12 proteins (50% identity clustering) identified from metagenomic data. External ring annotation corresponds to protein sequence length, assigned taxonomy at the domain and phylum level (20 most abundant phyla), and presence of Cas1/Cas2 proteins or a CRISPR array within 3000 bp flanking the TnpB locus. Protein length shows a bimodal distribution, with a peak around 400 aa, corresponding to TnpB and small Cas12 variants, and a second peak around 1,300 aa, corresponding to large Cas12 variants. TnpB and Cas12 orthologs are present in both Bacteria and Archaea, with the majority originating from Firmicutes, Proteobacteria, Actinobacteria, Euryarchaeota, and Cyanobacteria. Notably, closely related variants are often found in distantly related species (i.e., in different phyla or domains), likely due to horizontal transfer of the mobile elements that encode them. Node color indicates whether identified proteins can be classified as TnpB (no Cas1/2 or CRISPR array), known Cas12 (subtypes annotated by CRISPRCasTyper) or unknown Cas12 (not annotated by CRISPRCasTyper but having Cas1/2 or a CRISPR array). Known Cas12 subtypes are annotated along the tree, as well as TnpB variants active in human cells identified here and in previous studies ^1–3^.

**Extended Data Fig. 2.**
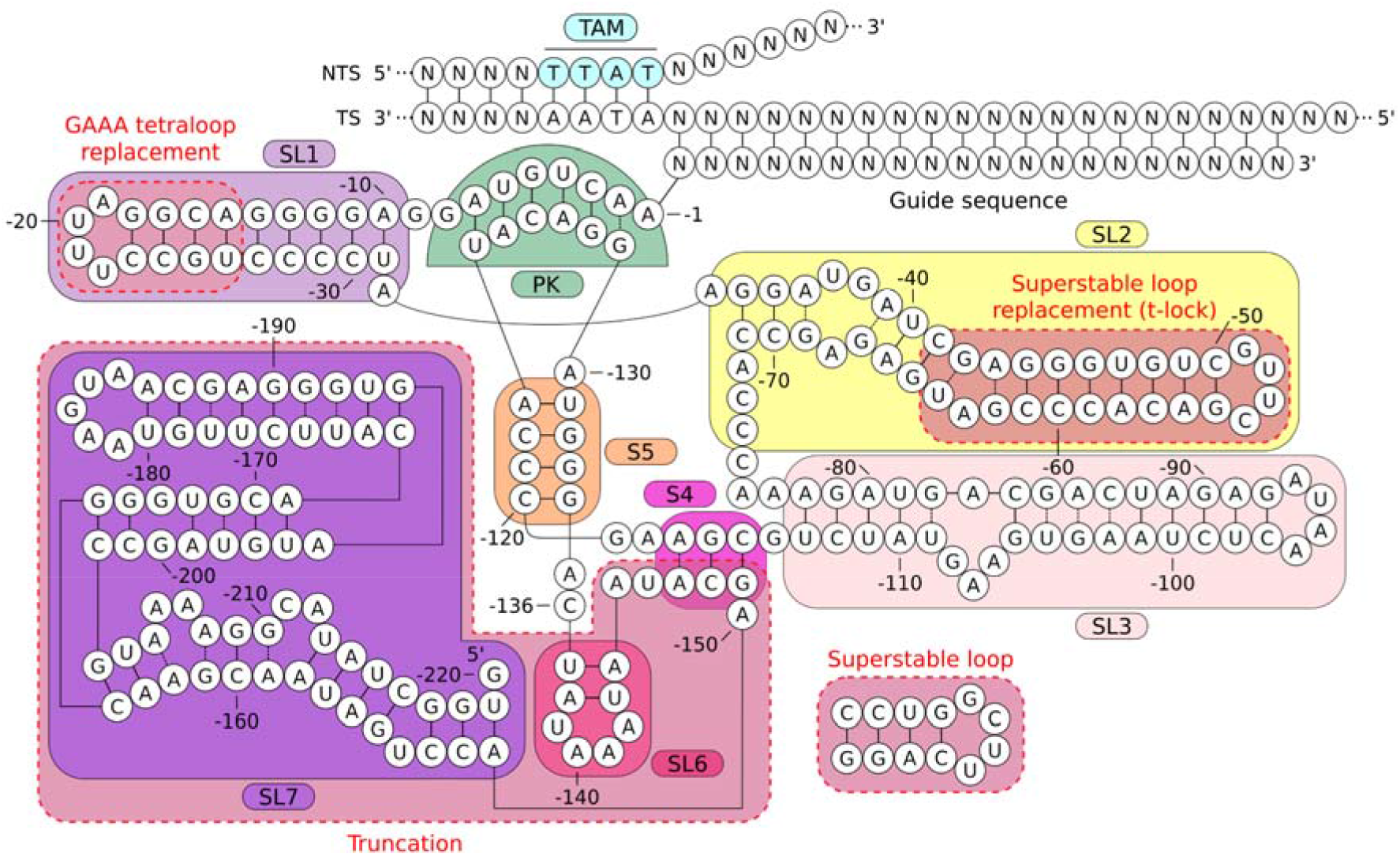
ISPmu1 ⍵RNA structural engineering. Predicted ISPmu1 ⍵RNA structure generated by combining secondary structure prediction (mfold) and tertiary structure prediction (AlphaFold3). A pseudoknot (PK), five stem loops and two internal stems were identified. The full-length ⍵RNA (220 bp) was first truncated to remove SL6 and SL7, generating the shorter 136 bp ⍵RNA with enhanced activity. ⍵RNA-v12 shortens stem loop 1 by replacing its distal end with a GAAA tetraloop, whereas ⍵RNA-v14 shortens and stabilizes stem loop 2 by replacing its distal region with a superstable loop (T-lock architecture) ^4^.

**Extended Data Fig. 3.**
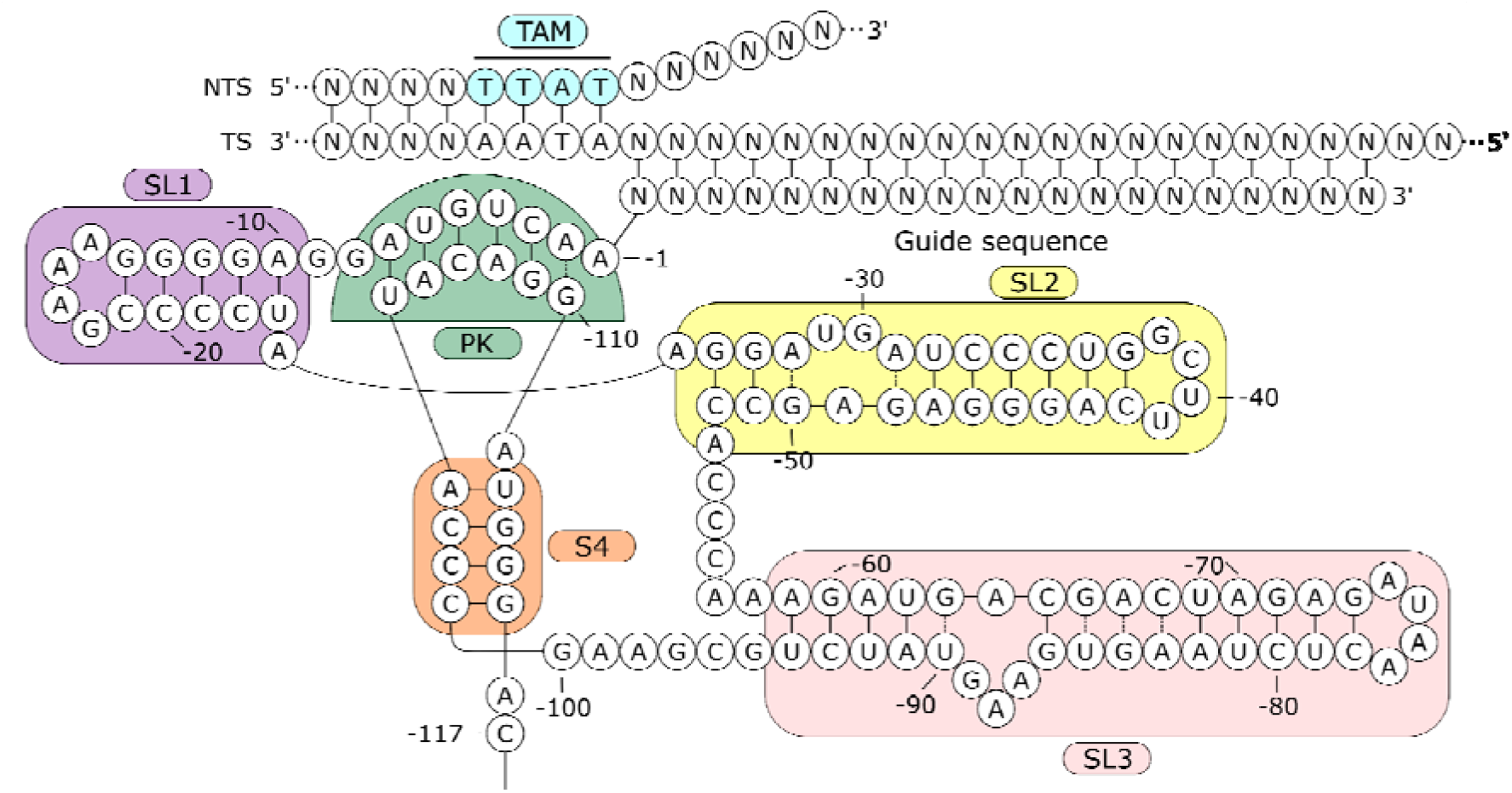
ISPmu1 ⍵RNA-v12+14 final structure. Schematic representation of ISPmu1 ⍵RNA-v12+14 structure, obtained by secondary structure (mfold) and tertiary structure prediction (AlphaFold3). The ⍵RNA-v12+14 construct was generated by combining the structural modifications from v12 and v14.

**Extended Data Fig. 4.**
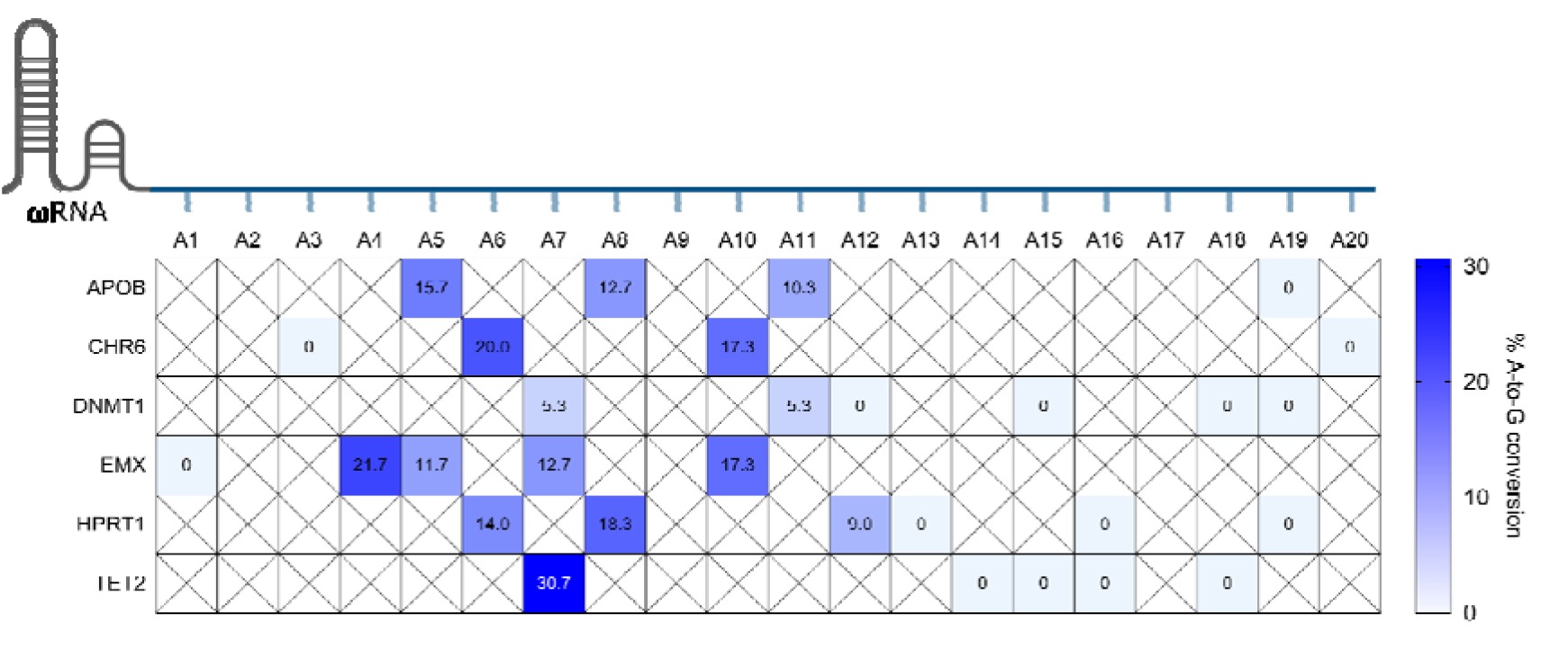
dEn-ISPmu1-ABE8e base editing efficiency. Adenine base editing at six individual human target sites. Mean A-to-G conversion (n=3) is indicated in the grid cells. Nucleotide positions along the guide (AN) are indicated at the top of the graph.

**Extended Data Fig. 5.**
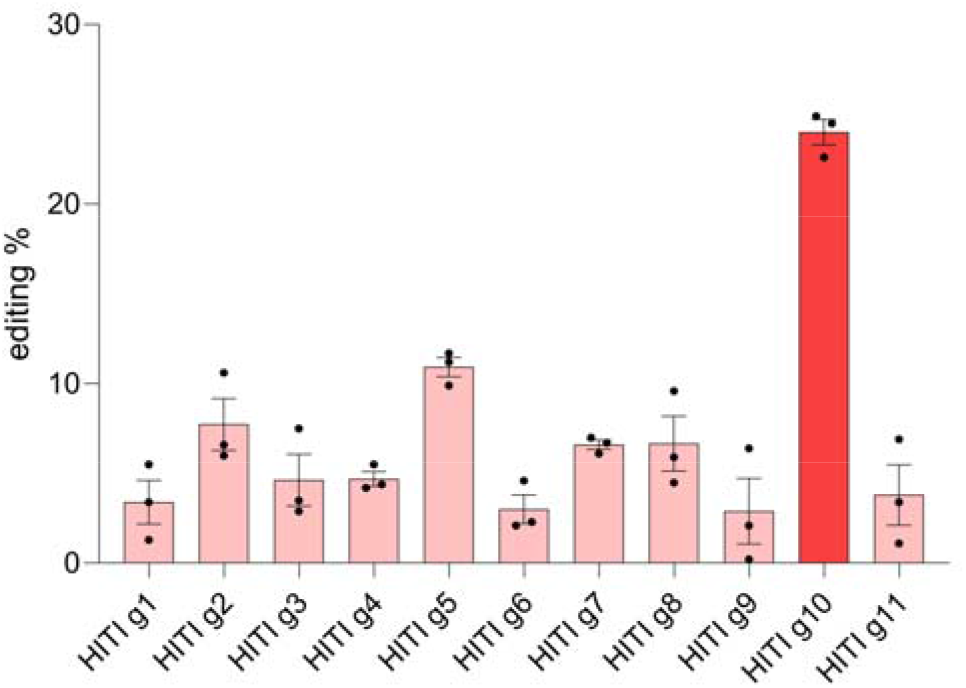
Guide screening on the mouse *Albumin* locus. Editing efficiency of 11 candidate guides spanning introns 13 and 14 of the *Alb* gene. The highlighted bar indicates the top-performing guide selected for subsequent HITI applications. Data represent mean ± SEM (n = 3 independent replicates).

**Extended Data Fig. 6.**
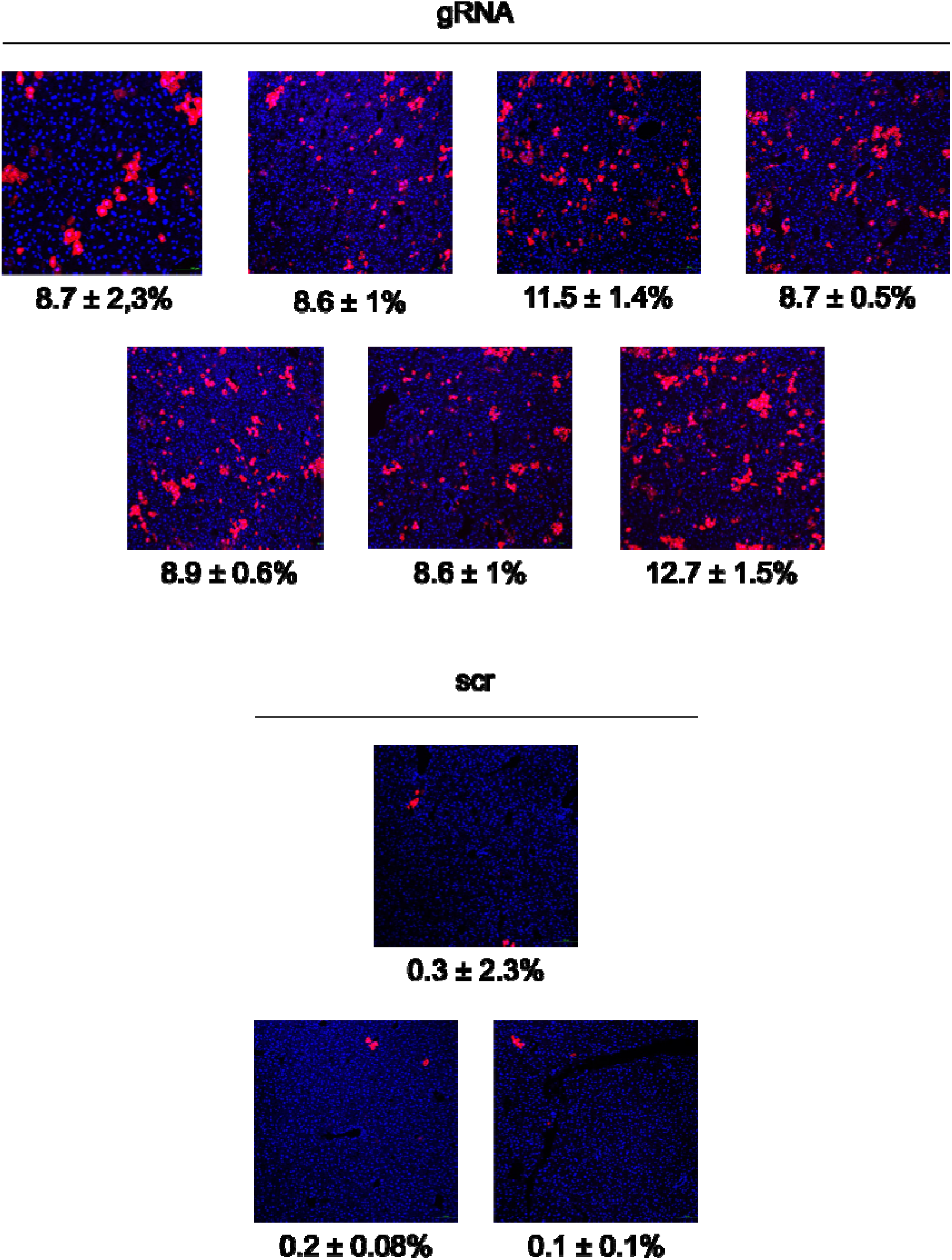
AAV-HITI-mediated transgene expression in neonatal liver. Fluorescence microscopy images of liver cryo-sections from wild-type mice injected with AAV8-DsRed transgene (gRNA, n= 7 or scrRNA, n = 3) at the total dose of 1.2 × 10^14 GC/kg. HITI efficiency ± SD are reported below the images.

