## Supplementary file for "Single-vector AAV delivery of En-ISPmu1 and donor DNA for *in vivo* targeted genome integration"

**SUPPLEMENTARY INFORMATION**

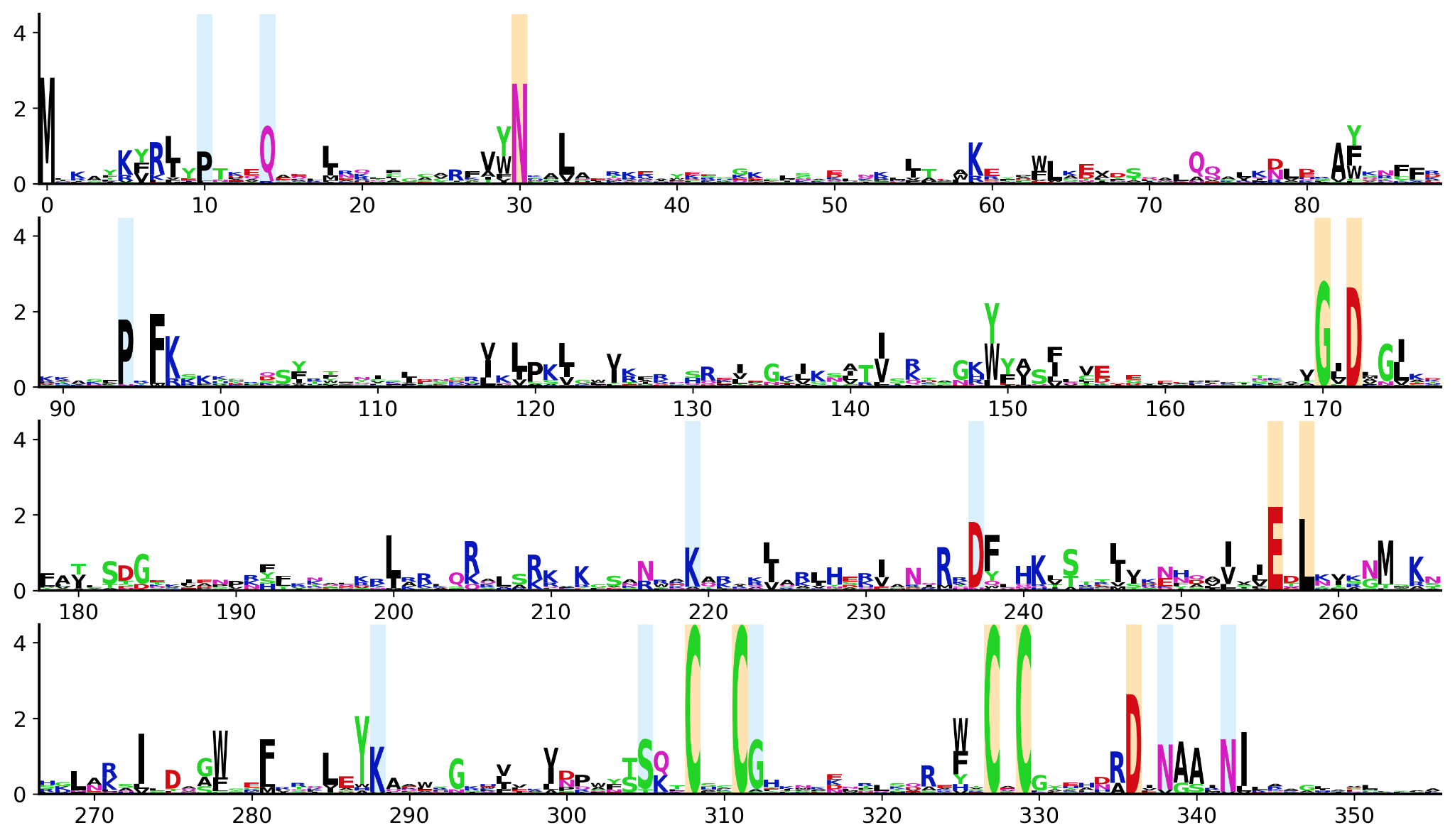

**Supplementary Figure 1. TnpB sequence logo.**

TnpB profile HMM logo generated from the multiple sequence alignment of TnpB orthologs showing detectable activity in *E. coli* in (Xiang et al. 2023). Residues conserved in >90% of active variants are highlighted in orange, if previously identified in (Xiang et al. 2023), or in blue, if identified here.

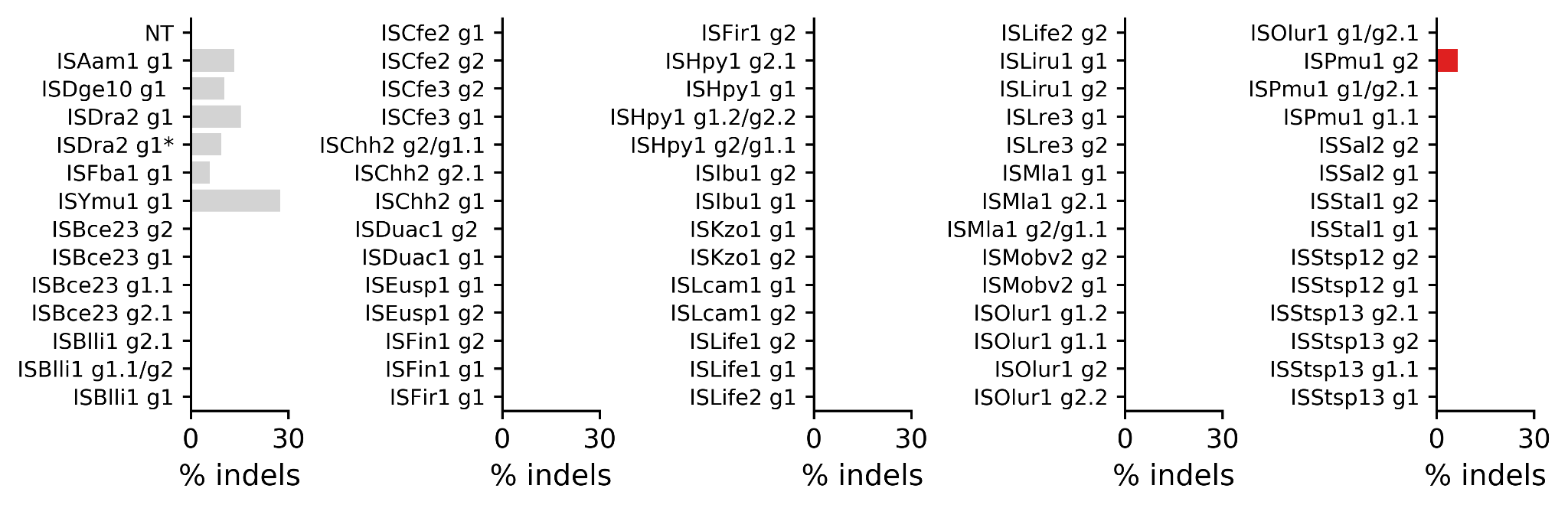

**Supplementary Figure 2. TnpB screening in HEK293T cells.**

Editing activity of 25 selected TnpB variants in HEK293T cells, measured by indel formation at the EMX1 locus. Positive controls are ISAam1, ISDge10, ISDra2 (with full length ⍵RNA g1 and truncated ⍵RNA g1*), ISFba1 and ISYmu1. Multiple guides were tested for each variant, corresponding to different TAM sequence predictions.

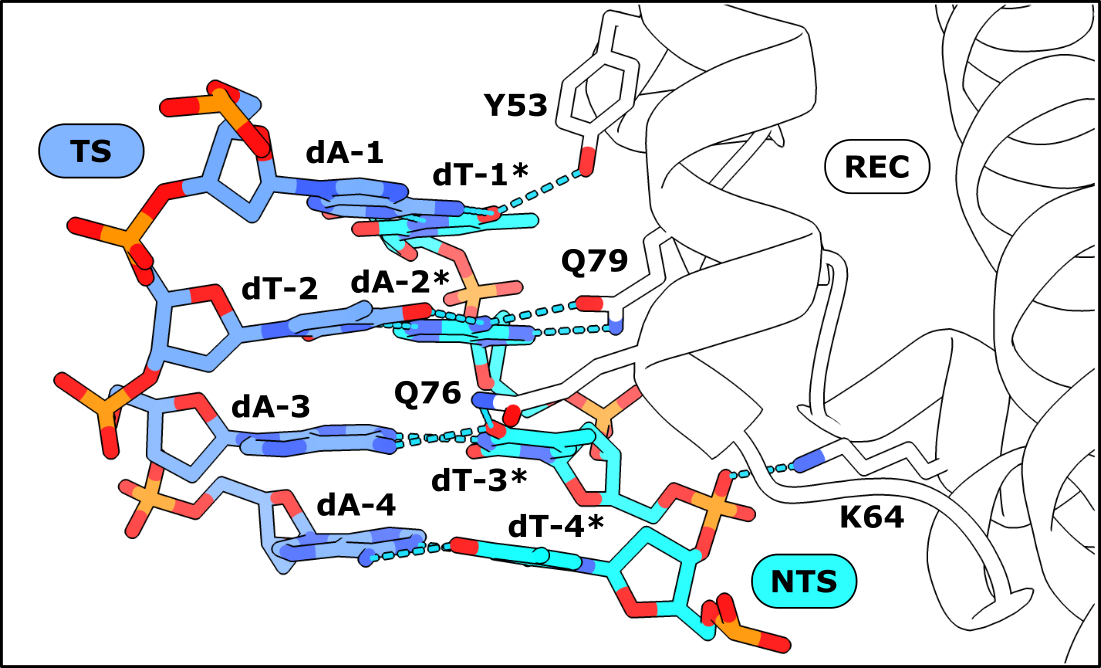

**Supplementary Figure 3. AlphaFold3 prediction of TAM recognition by the REC domain.** Residues Y53, Q79 and Q76 form hydrogen bonds with non-target strand (NTS) residues dT-1*, dA-2* and dT-3*, respectively, while K64 interacts with the backbone phosphate group between dT-3* and dT-4*.

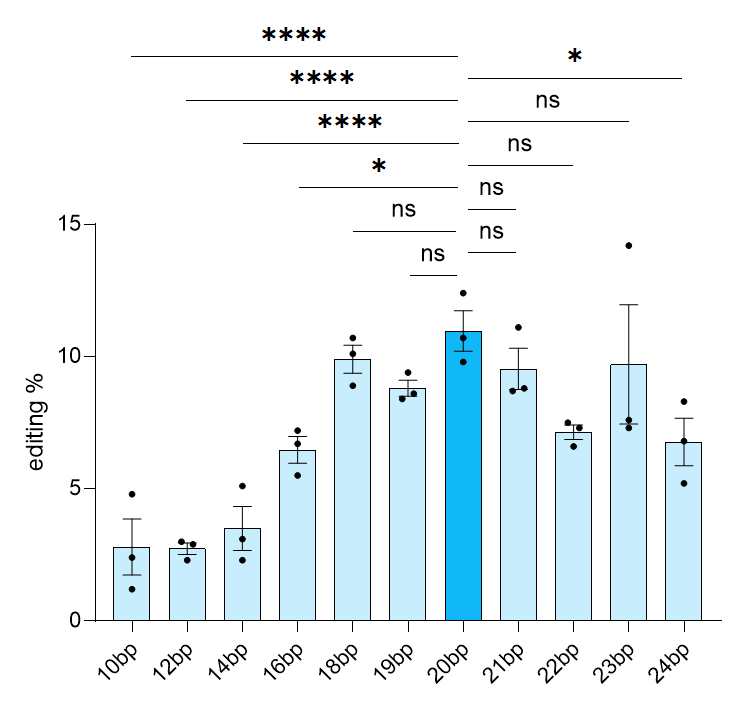

**Supplementary Figure 4. Guide length screening on *EMX1*.**

Editing efficiencies of ISPmu1 ⍵RNA v12+14 with varying spacer lengths (ranging from 10 to 24 nucleotides) targeting the EMX1 locus. The highlighted bar indicates the optimal guide length selected for subsequent experiments. Data represent mean ± SEM (n=3 independent replicates). Comparisons were performed using a one-way ANOVA followed by Dunnett's multiple comparisons test against the reference.

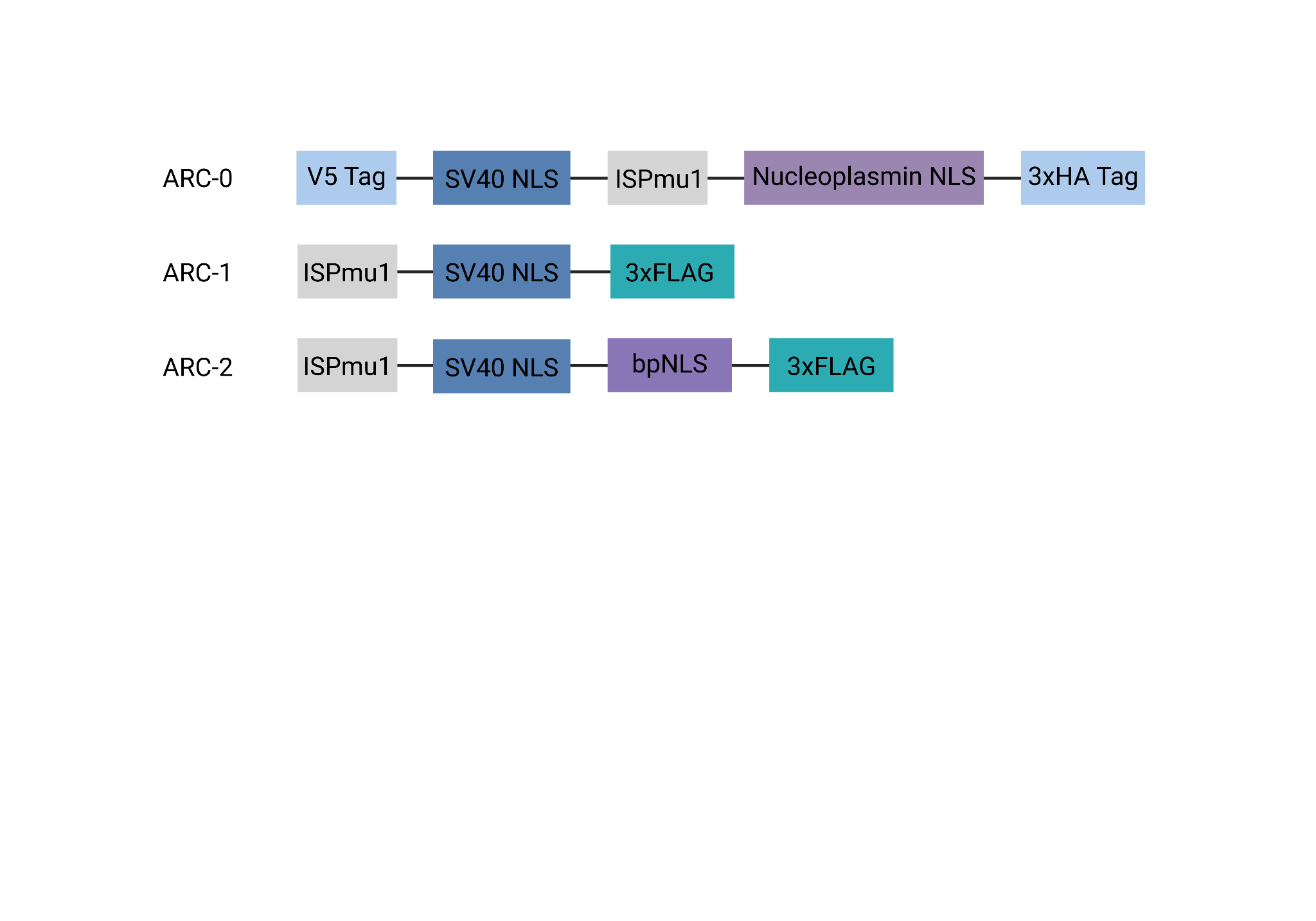

**Supplementary Figure 5. Schematic representation of tested ISPmu1 architectures.** Structural layouts of the evaluated ISPmu1 constructs, detailing the relative positions of the protein, nuclear localization sequences (NLS) and tags.

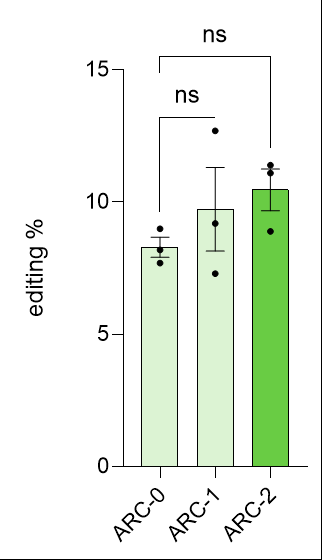

**Supplementary Figure 6. Editing efficiency of ISPmu1 architectures on *EMX1*.**

Genome editing efficiencies across the distinct ISPmu1 configurations. The highlighted bar indicates the optimal architecture selected for subsequent experiments. Data represent mean ± SEM (n=3 independent replicates). Comparisons were performed using an unpaired t-test with Welch’s correction (ns p ≥ 0.05).

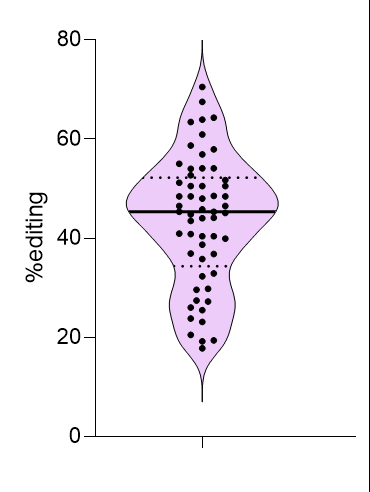

**Supplementary Figure 7. Summary of editing efficiencies across human endogenous loci.**

Violin plot displaying the distribution of indel frequencies achieved by En-ISPmu1 across a panel of human endogenous sites (loci=12; n=3 independent replicates per locus). The thick line indicates the average editing, dotted lines denote the 25th and 75th percentiles.

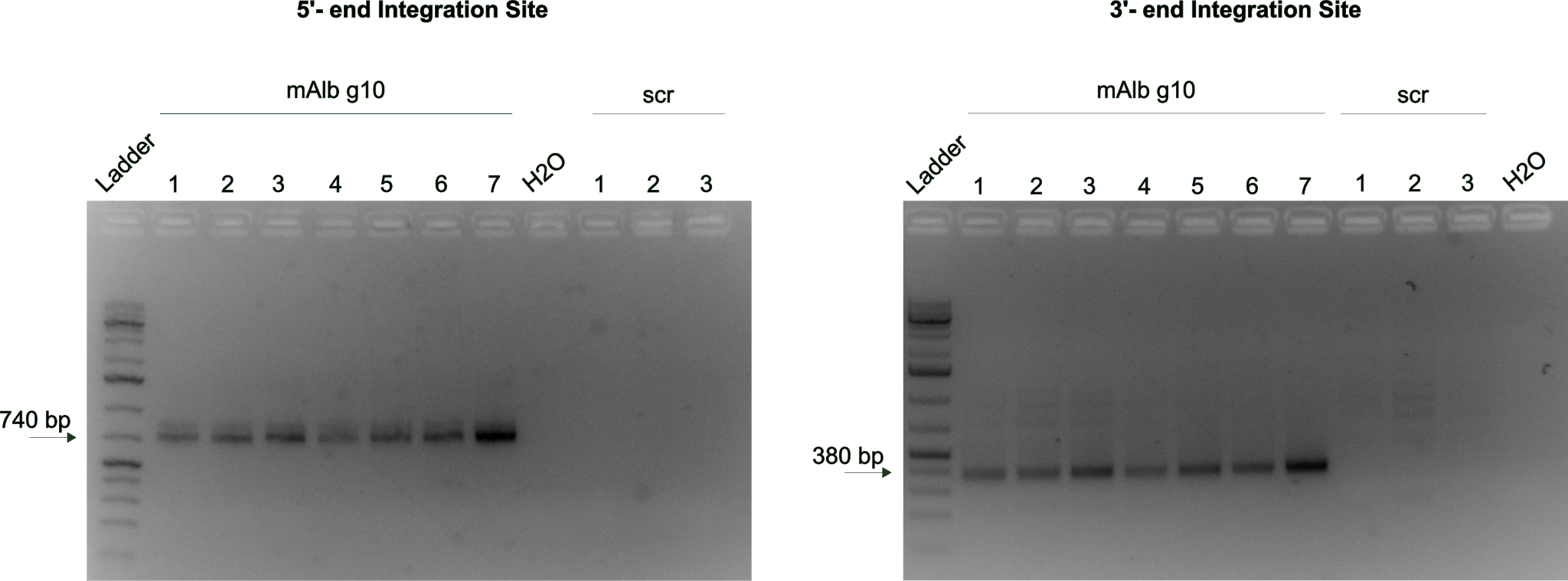

**Supplementary Figure 8. PCR verification of integration junctions.**

Agarose gels displaying expected in-out PCR amplicons to validate site-specific genomic integration at the target *Albumin* locus. Lanes correspond to individual mice (mAlb-g10 n=7; scr n=3), with Gene Ruler 1kb Plus ladder used as the molecular weight standard.
